# Characterizing the Human Heart Rate Circadian Pacemaker through Widely Available Wearable Devices

**DOI:** 10.1101/2020.03.12.988899

**Authors:** Clark Bowman, Yitong Huang, Olivia J Walch, Yu Fang, Elena Frank, Cathy Goldstein, Srijan Sen, Daniel B. Forger

## Abstract

The effects of real-world stimuli and how they vary across individuals remains largely unknown for lack of a continuous real-world circadian marker. Here, we show how an underlying circadian rhythm in heart rate (CRHR) can be extracted from several widely used wearables. Analysis of over 130,000 days of data from medical interns on rotating shifts shows how CRHR dynamics are distinct from those of sleep-wake timing and vary greatly among individuals. Analysis of circadian timekeeping in travelers in 5 continents shows how the circadian timekeeping more carefully controls wake time rather than sleep time. We determine a personalized phase response curve (PRC) of CRHR to activity for each individual, representing the first passive and personalized determination of how human circadian timekeeping continuously changes due to real-world stimuli. These results collectively establish CRHR as a practical method to study circadian rhythms in the real world.

**eTOC Blurb:** We show how the circadian rhythm in heart rate can be extracted from real world data collected by wearables. Studying data from a large cohort of medical interns working on shifts, we find very interesting dynamics of this circadian rhythm that are independent of the acute effects of activity or sleep-wake timing. These techniques can also determine personalized parameters of circadian timekeeping.

## Introduction

The human circadian rhythm, the internal clock synchronizing physiological functions with the daily cycle, is known to have wide-ranging connections to human health. While markers of this clock such as melatonin can easily be measured with clinical procedures (e.g., blood or saliva samples), the vast body of research on circadian rhythms cannot be translated to population-scale applications without a passive way to assess circadian phase in the real world. From optimally timing chemotherapy to better designing shift schedules to maximize the performance and health of shift workers, the benefits of such a method would be enormous (Arendt, 2010; Crowley et al., 2004; Sahar and Sassone-Corsi, 2009).

The recent popularity of wearable devices provides a new opportunity for real-world circadian assessment. Many widely used wearables continuously measure heart rate, which could theoretically measure an underlying circadian rhythm in heart rate (CRHR); sleep pioneers such as Kleitman and Timmerman recognized heart rate as a circadian marker as early as the 1950’s, but debated its usefulness since its underlying rhythm was masked by the effects of sleep and activity (Kleitman and Kleitman, 1953; Timmerman et al., 1959). As a result, CRHR has to date been practical only for measuring circadian phase in “constant routines,” wherein sleep and activity are carefully controlled. Though constant routines are useful for clinical studies, their cost and invasiveness preclude their use in large-scale applications.

Here, we propose a probabilistic method for extracting CRHR from real-world wearable data. As in existing methods for studying core body temperature data, we allow for temporal dynamics in HR and remove the effects of activity (Brown and Czeisler, 1992). To avoid the masking effect of sleep noted by Kleitman et al., we use the most conservative method possible: discarding all data during episodes of sleep. Both because of the removal of sleep data and because wearables are often removed for charging, the resulting data will have large gaps; such gaps are known to bias many current mathematical techniques for phase estimation (Huang et al., 2019a). We therefore employ a recent approach of Huang et al. to avoid this bias. We show that: 1) the rhythms we extract match, on average, the rhythms measured in constant routine; 2) our analysis of heart rate over the course of a day matches many known properties of heart rate; 3) the CRHR is shifted by, but does not directly track, cues such as light and sleep, similar to other measurements of the human circadian pacemaker; 4) our methods perform similarly for the three different wearable devices we test; and 5) CRHR validates against the rhythms measured by dim light melatonin onset (DLMO), the current gold standard in circadian measurement. We conclude by demonstrating potential uses of CRHR as a circadian marker in understanding inter-individual differences in circadian timekeeping using over 130,000 days of real-world data.

## Results

### Estimating the circadian rhythm in heart rate (CRHR)

We first introduce our method for extracting circadian information from wearable HR data. As a baseline, we assume a 24-hour circadian rhythm in HR. We assume HR increases from this circadian baseline proportionate to activity, matching existing data (Scheer et al., 2010). A simple linear model, i.e., each step increasing cardiac demand by a fixed amount, was found to work as well as more complex models - we rarely observed cases of heart rate approaching a maximal value in our data set. Since sleep also masks rhythmicity in heart rate, we removed all data obtained when the device reported the user to be asleep.

HR measurements are intrinsically noisy on wearable devices. Moreover, there are many processes and factors other than steps which affect heart rate. We therefore assume a simple autoregressive (AR) model, which includes both measurement noise and short-term correlated noise, for the error term. Due to the assumption of correlated errors, we chose a likelihood-based approach to fit model parameters. The likelihood approach provides error estimates and is not affected by large gaps in data, such as when devices are charged, which can bias other approaches such as least-squares (Huang et al., 2019a). It also allows us to easily impose continuity: when uncertainty is low, circadian phase is not likely to shift more than one or two hours in a given day. (When uncertainty is high, the estimated phase can still jump several hours between days.) Figure 1A shows the average fit for a real individual from our data set.

**Figure 1:**
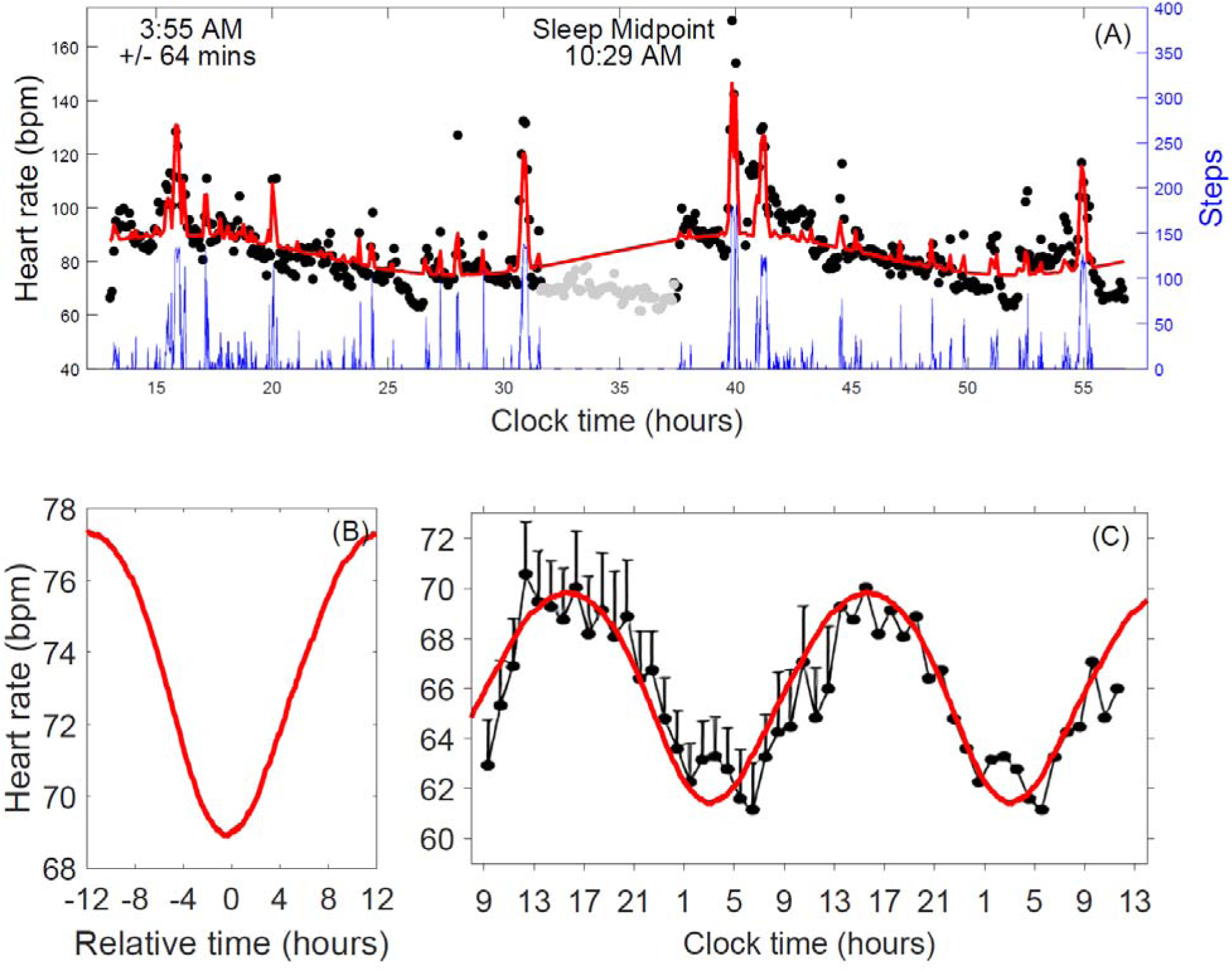
Measurement and validation of circadian rhythm in heart rate (CRHR) (A) Model fit (red curve) for two days of real data on heart rate (black dots) and activity (blue curve). Days are separated by a period of sleep (gray dots) centered at 10:29 AM; the HR clock phase minimum at 3:55 AM (+/- 64 mins) indicates desynchrony (CRHR phase is advanced compared to the sleep-wake cycle). (B) Average human background heart rate estimated by our model from 136,789 days of Fitbit data. Background heart rate was obtained by taking heart rate measurements during periods of wakefulness and subtracting out the estimated linear effect of activity, then shifting by estimated CRHR phase. On average, background heart rate was found to oscillate by roughly 8.32 bpm throughout the day. (C) Comparison of measured background rhythm in heart rate (red) with the rhythm observed by G. Vandewalle et al. in constant routine (black) (Vandewalle et al., 2007). Due to natural variability in resting heart rate and lack of a controlled setting, the average heart rate in our data set (73.52 bpm, n = 927) was higher than the average observed in the constant routine study (∼66 bpm, n = 8) and was shifted downward to match. Visually, our model recovers the amplitude and phase of the circadian oscillation in heart rate to good accuracy.

To demonstrate that our model captures the true underlying rhythmicity in heart rate, we fitted heart rate models for 136,789 days of Fitbit data from the Intern Health Study, an annual cohort study that utilizes physician training as a model to understand how various forms of stress (including shifting schedules) increases risk for disease (Sen et al., 2010). While we largely focus on estimating the phase of circadian heart rate rhythmicity, our model also fits its amplitude and vertical offset; Figure 1B shows the average measured background oscillation in heart rate over the entire data set. Vandewalle et al. measured a similar curve in experiment under “constant routine”, wherein activity and sleep were carefully controlled (Vandewalle et al., 2007). Figure 1C shows our measurements to match those of Vandewalle et al. to high accuracy: our approach accurately captures the known background oscillation in heart rate from real-world, noisy Fitbit data.

### Dynamics of real-world CRHR

We next examine the dynamics of CRHR as a circadian marker in real-world data; several applications of heart rate phase measurement are collected in Figure 2.

**Figure 2.**
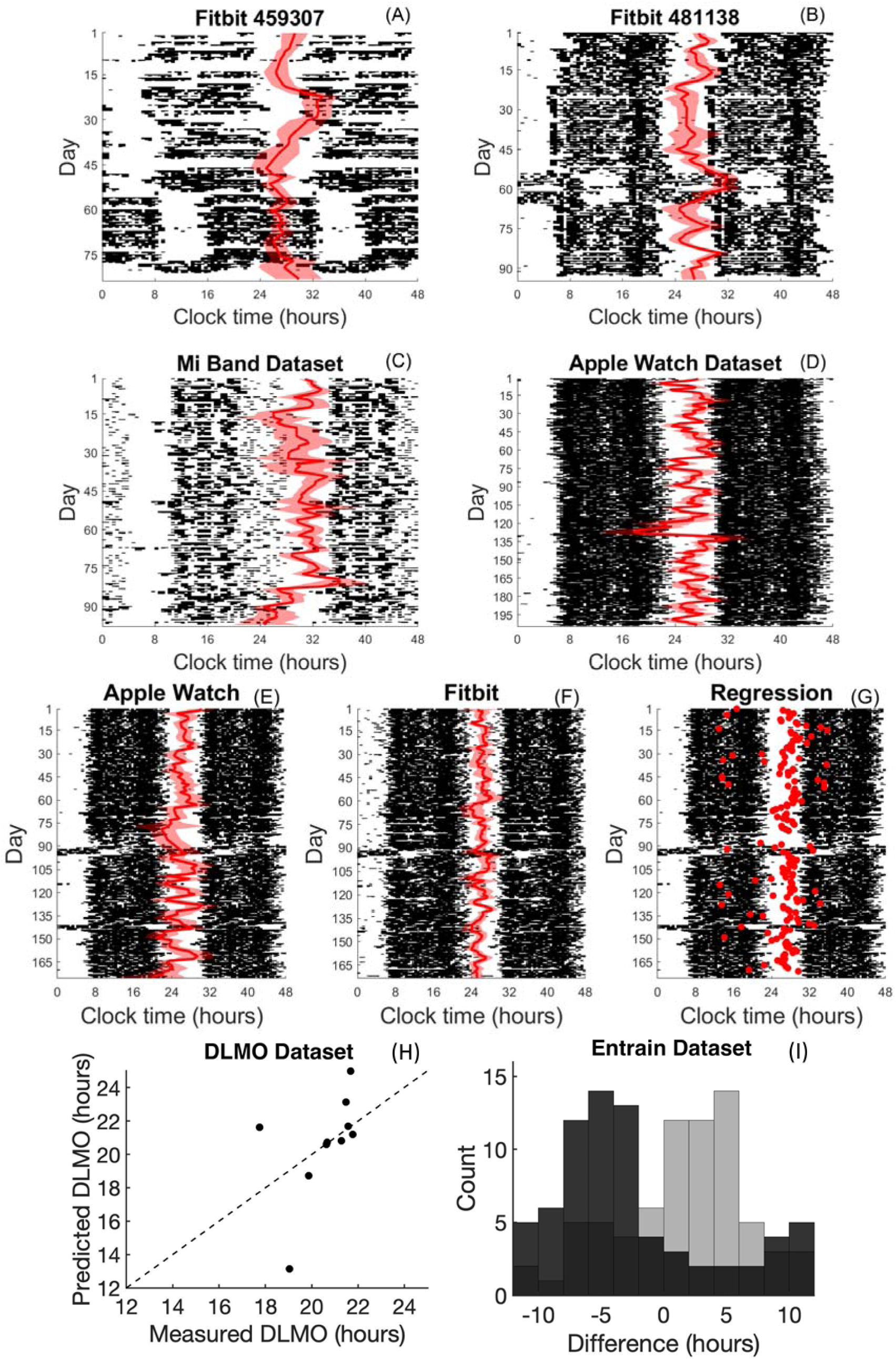
Dynamics of CRHR in real-world data. (A, B) Actograms for two subjects in the Intern Health Study generated from Fitbit data. Estimated CRHR phase (red, with 80% confidence bands) is overlaid on daily activity patterns (black). Subject A maintains a consistent CRHR throughout a shifted sleep schedule, while Subject B gradually adjusts. (C) Estimated CRHR phases for the Mi Band data set. The lower accuracy of Mi Band HR measurements results in wider confidence bands. (D) Estimated CRHR phases for the Apple Watch data set. Despite discarding data obtained during sleep, estimates fluctuate around sleep midpoint. (E, F) Comparison of estimated CRHR phase for an individual wearing both a Fitbit and an Apple Watch simultaneously; results are strongly correlated (p = 0.00096), showing that both devices measure the same underlying rhythm. (G) Estimated CRHR phase (red circles) for the same Fitbit/Apple Watch individual using orthogonalized least-squares regression, the simple approximation which can run on smartphones. (H) Estimating DLMO from the heart rate model for 10 subjects. Estimates largely agree with measured DLMO (7 subjects within ∼2 hours). For individuals with typical sleep schedules, the SCN and HR clocks appear to be synchronized. (I) Comparison between users’ self-reported sleep schedules and estimated CRHR phase for 72 individuals in the *Entrain* data set (Walch et al., 2016). Differences between wake time and CRHR phase are shown in black and differences between time to bed and CRHR phase in gray; wake time is a better predictor of CRHR phase than is time to bed.

We first use our model to extract phase for a representative selection of individuals. To show robustness across devices, we supplement the 136,789 days of intern Fitbit data with additional data from an Apple Watch (1,071 days) and a Mi Band (189 days) measured from two authors of this study. Figures 2A-2D show actograms for two interns from the Intern Health Study (A, B), the Mi Band author (C), and the Apple Watch author (D). Daily activity patterns, in black, show individuals A and B to transition through a period of shiftwork, while individuals C and D maintain consistent activity patterns. When daily routines are consistent, CRHR phase estimates (red line) and 80% confidence bands (wide red region) largely track with sleep (overnight gaps in activity).

Conversely, major disruptions in daily routine yield a range of different behaviors in CRHR. Intern A (Fitbit 459307) provides a useful example: in days 1 - 50, this user’s activity pattern slowly adjusts to a later schedule, and the clock in HR seems to adjust quickly (and perhaps overadjusts). A new work shift begins around day 55, yielding a dramatic 7-hour shift in activity pattern; however, the HR clock does not shift, instead following other consistent cues rather than sleep timing - perhaps activity, light exposure, or meals. Intern B (Fitbit 481138) begins a similar shift around day 50, but in this case we see CRHR slowly entraining to the new schedule. Shift workers use different strategies to adjust to new schedules; this is, to our knowledge, the first large-scale continuous measurement of a circadian marker during these adjustment periods.

Figures 2C-D show data from the Mi Band and the Apple Watch, with estimated CRHR phase roughly synchronized with a constant sleep routine for several consecutive months in individuals without shiftwork. As a more direct comparison of devices, one author of this study wore both a Fitbit and an Apple Watch for a period of six months. Figures 2E and 2F show that the resulting heart rate circadian phase estimates were strongly correlated (p = 0.00096). Note that both Fitbit and Apple Watch data suggest that, for this individual, CRHR remained consistent through a period of travel around day 90 which resulted in a major shift in activity patterns. Figure 2G estimates CRHR phase for the same subject using a slightly modified approach based on regression which can feasibly run on a smartphone. We use the orthogonalized approach of Huang et al. rather than ordinary least squares to avoid bias when there exist large gaps in measurements (e.g., during sleep or when the device is charged) (Huang et al., 2019a). For most days, the resulting phase estimates agree well with the likelihood-based estimates shown in Figures 2E and 2F.

We next validate our heart rate phase estimates against DLMO, widely considered the gold standard in circadian measurement. We recruited subjects for laboratory DLMO measurements; each was given an Apple Watch to wear in the preceding weeks (Walch et al., 2019). While use of the watch varied, many individuals wore the watch for most of each day: of the 23 subjects who exhibited melatonin onset during this sleep study (i.e., had a valid DLMO measurement), 10 had at least seven days of consecutive heart rate and activity data.

To predict DLMO using heart rate measurements, we shifted CRHR phases by a fixed amount to account for the phase difference between DLMO and circadian minimum; Figure 2H compares these estimates with measured DLMO. The mean absolute difference between the predicted and measured DLMO phases was 1.717 hours, with 7 of 10 subjects agreeing to within 2 hours. For individuals on normal sleep schedules, the suprachiasmatic nucleus (SCN) clock, which regulates DLMO, appears to be synchronized with the HR clock.

Finally, we test a result of Walch et al. (Walch et al., 2016), who suggested based on survey data from the mobile app *Entrain* that wake time correlates better with circadian phase than does time to bed. Within the *Entrain* data, 72 individuals, spanning 5 continents, anonymously submitted through the app enough heart rate and activity data to allow prediction of CRHR phase. We observed the same effect as Walch et al. (Figure 2I), i.e., differences between CRHR phase and wake time vary less (σ = 1.15 hours) than differences between CRHR phase and time to bed (σ = 1.21 hours), despite the strong correlation between time to bed and wake time via average sleep duration. This serves as a verification of the central hypothesis of Walch et al.: wake time is more closely controlled by the circadian pacemaker than is the beginning of sleep. This is one of many key properties shared by both CRHR and the SCN clock.

### Relationship between CRHR and sleep

The results collected in Figure 2 illustrate the general principle we see in the data set at large: while CRHR tends to synchronize with sleep-wake patterns, it does act independently and can be driven significantly out-of-phase from the sleep cycle. Since all data obtained during sleep were discarded, this result cannot be affected by the masking effect of sleep on heart rate. An analysis of the residual correlations, shown in Figures 3A – 3D, provides further evidence that these processes are independent: clocks which were estimated as being out of phase with the sleep-wake cycle tend to stay out of phase, synchronizing only gradually over several days (or sometimes not at all).

**Figure 3.**
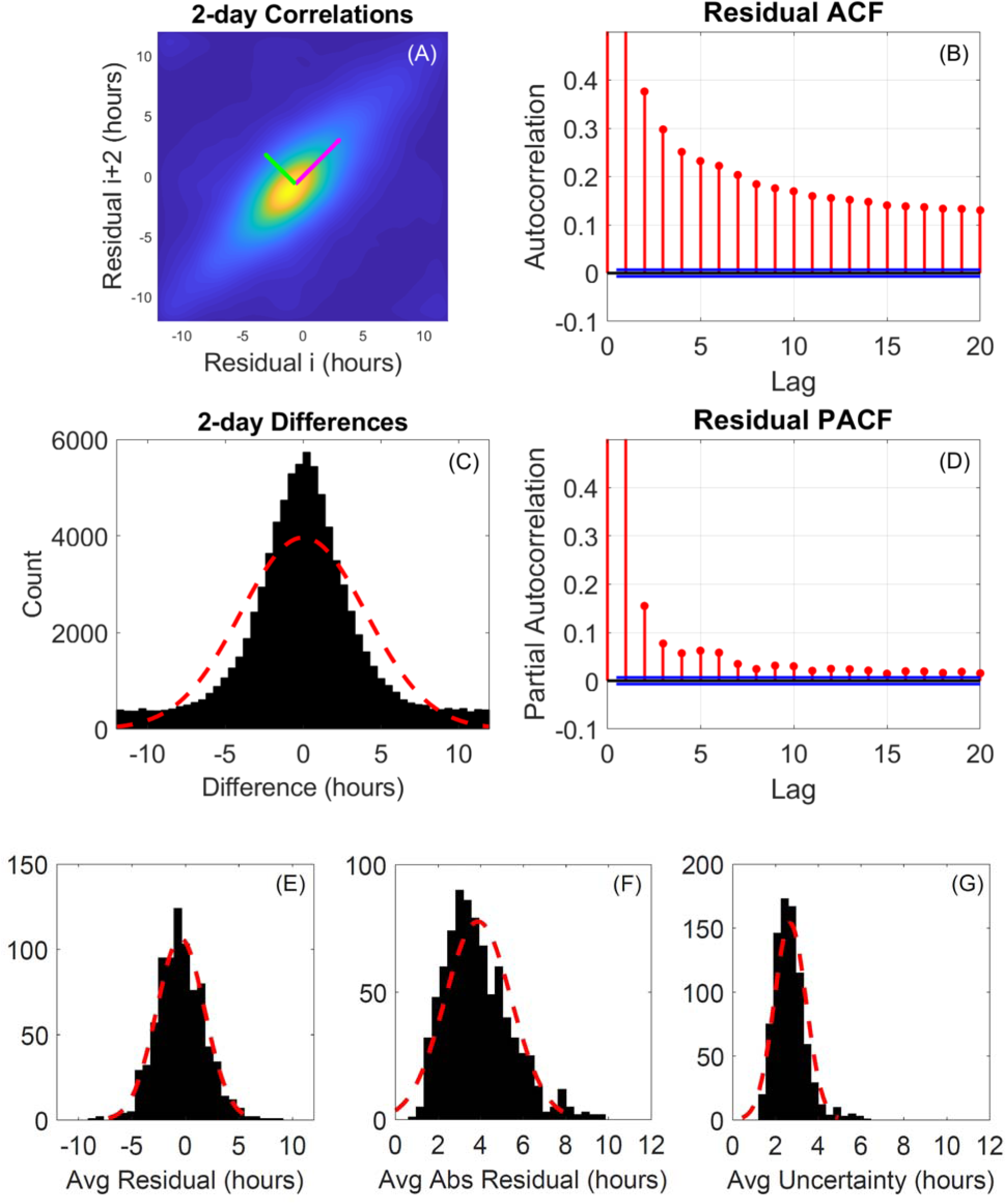
Analysis of phase difference (residual) between midsleep and CRHR. (A) Comparison of residuals (difference between CRHR phase and sleep midpoint) separated by two days. Pink arrow along the major diagonal is the direction of largest variation: when the HR clock is out of phase with the sleep-wake cycle, it is likely to be similarly out of phase in two days. (B) Autocorrelation function (ACF) of residuals. Blue lines are p = 0.05 assuming independence. Residuals are highly correlated over consecutive days. (C) Histogram of changes in residual over two days (μ = 0.03, σ = 4.03 hours). Accounting for uncertainty, residuals rarely change by more than 2-3 hours over a two-day period. (D) Partial autocorrelation function (PACF) of residuals. After subtracting out short-term correlations, the HR clock synchronizes with the sleep-wake cycle on the timescale of 1-2 weeks. (E) Histogram of average residual across all subjects (μ = −0.45, σ = 2.25 hours). (F) Histogram of average absolute residual across all subjects (μ = 3.88, σ = 1.56 hours), which includes errors due to uncertainty and shifting sleep schedules. (G) Histogram of average uncertainty across all subjects (μ = 2.67, σ = 0.74 hours). For most subjects, phase is usually estimated to within 2-3 hours.

Compared to DLMO, a major advantage of a HR-based marker for circadian phase is the amount of available data: the Intern Health data set contains over 130,000 days of minute-by-minute Fitbit data from more than 900 individuals working shifts across the United States. We can therefore provide a wide-ranging characterization of the human circadian rhythm in heart rate (Figures 3E-3G). The average phase difference between sleep midpoint and CRHR phase, which we refer to as the residual, has a larger range than one would expect from DLMO. For most individuals, this is the average of hundreds of days of data, meaning they maintain a stable phase difference with sleep.

Figure 3E shows that some individuals sleep at earlier phases of their CRHR than do others. Our measurements of CRHR suggest that sleep timing alone cannot determine whether a person’s circadian rhythms are advanced or delayed. The average absolute residual, shown in Figure 3F, corroborates this result, though it also incorporates errors due to uncertainty and shifts in the sleep schedule (which may temporarily drive the HR clock out of phase with sleep, even if they are usually synchronous). The average uncertainty in our phase estimate is shown in Figure 3G. For most individuals, phase is usually estimated to within +/-2-3 hours.

### A personalized phase response curve of CRHR

We hypothesized that activity (and related factors, such as light exposure, which correlate with measured activity) would be a suitable proxy for the signals which entrain the HR clock; recent modeling work suggests that activity measurements can outperform even light measurements in predicting DLMO (Huang et al., 2019b). To characterize this effect, we calculated a phase response curve (PRC) for each individual (see Methods) to activity, which is measured by the devices via “steps”. In brief, we calculate statistically how much activity at different times of the day phase shifts CRHR; if the rhythms in heart rate were not shifted by activity, we would expect to see only random noise. In contrast, we found a robust PRC for basically all individuals (Figure 4).

**Figure 4.**
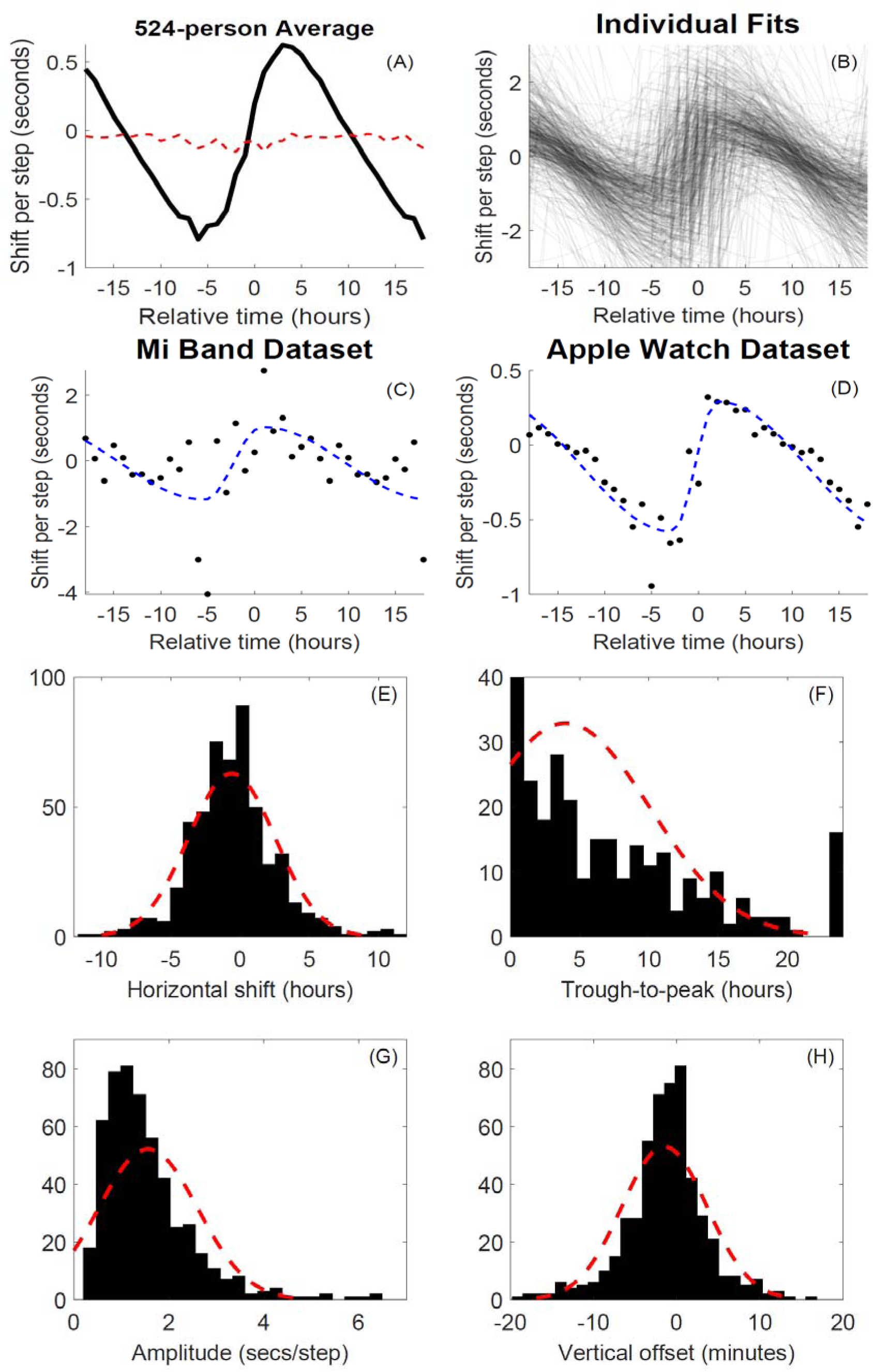
Individualized phase response curves of CRHR. (A) Average phase response curve (black) for all subjects with at least 50 days of data. Activity shifts the HR clock, but the effect varies strongly with time (relative to the current phase). When hours of activity are randomly shuffled, the relationship vanishes (dashed red line). (B) Cloud of individual phase response curves for 524 Intern Health Study subjects. (C) Raw PRC data (black, 1 hr bins) with parametrized fit (dashed blue) for the Mi Band data set (189 days). (D) Raw PRC data with parametrized fit for the Apple Watch data set (1,071 days). (E) Histogram of horizontal offsets for individual PRCs (μ = −0.61, σ = 3.14 hours; gaussian fit shown with dashed red line). Zero means the curve crosses from negative to positive exactly when baseline HR is at its lowest. (F) Histogram of trough-to-peak values (time between minimum and maximum) for individual PRCs (μ = 3.97, σ = 6.06 hours). Many individuals did not have sufficient steps at all phases to resolve this parameter well; values at 0 (bar extends upward) and 24 are half-period fits with jump discontinuities. (G) Histogram of individual PRC amplitudes (μ = 1.56, σ = 1.05 seconds per step). (H) Histogram of individual PRC vertical offsets scaled by average steps (μ = −1.48, σ = 5.14 minutes); negative values should indicate clock periods longer than 24 hours (positive values, shorter).

As a first comparison, we calculated the average PRC for all individuals with at least 50 days of data (Figure 4A). The result was almost identical in shape to the phase response curve to light previously measured for humans (Khalsa et al., 2003), including a nonzero vertical offset which has previously been linked with the non-24-hour intrinsic period of the clock.

We were also able to generate individual phase response curves for each subject (Figures 4B-4D). Since many tens or hundreds of consecutive daily measurements of the clock are needed to generate these curves, this would be impractical using DLMO techniques. These PRCs are completely personalized - they do not use data from other subjects or from the average “human” clock - and yielded consistent shapes both across individuals and across devices.

Interpersonal variation is examined more closely in Figures 4E – 4H. The horizontal shift (Figure 4H) varied among individuals but was similar to the spread observed between sleep midpoint and HR clock phase (Figure 3E). Figure 4F shows the trough-to-peak time, which measures sinusoidality of the curves. Recent work using the Ott-Antonsen theory shows how this is related to the coupling of individual cellular oscillators within the SCN (Hannay et al., 2015). The amplitude of the phase response curve (Figure 4G) may reflect conditioning to activity; for example, athletes may have clocks which are less shifted by activity than sedentary individuals, as their metabolic demands are different. Of particular interest, Figure 4H shows the offset of the PRC, which should measure the differences between the intrinsic period of the circadian clock among different individuals. This gives an average period of the HR clock as 24.03 +/- 0.08 hours, in line with previous work showing that human circadian clocks are tightly regulated to be slightly longer than 24 hours (Czeisler et al., 1999; Hiddinga et al., 1997).

Other parameters of the CRHR PRC are less tightly controlled. For example, amplitude may vary based on the individual’s overall activity level. The wide range of these parameters suggests the existence of significant interpersonal variation in the dynamics of the CRHR clock. Other circadian mechanisms exhibit similar variation. As a recent example, melatonin suppression due to light exposure was found to vary significantly between subjects (Phillips et al., 2019). Some of the interpersonal differences we observe in overall PRC shape could be due to related effects which change the synchrony of intracellular oscillations (Hannay et al., 2015).

## Discussion

Here, we describe a method whereby circadian phase can be passively assessed using readily available measurements of heart rate. Our analysis shows that the 24-hour rhythmicity in heart rate reflects an underlying clock and that CRHR is coordinated by the same timekeeping system as is measured in DLMO. Future work should better delineate the differences between these two circadian measures; their misalignment might be an important marker of internal desynchrony.

Understanding heart rate is vitally important because it is a critical marker of increased risk of cardiovascular disease. Our work clarifies the expected circadian rhythms in heart rate, potentially leading to more accurate measures of what constitutes a normal or abnormal heart rate. Further work is also needed to pin down the physiology of CRHR. Much of it is under the control of the autonomic nervous system, which also controls melatonin release. Occam’s razor would suggest that the molecular clock in cells of the sinoatrial node would also regulate this rhythm. The circadian clock in cardiomyocyte has been shown to regulate aspects of cardiac physiology.

Our method is subject to some limitations that should be noted. The effects of caffeine, psychological stress, and pharmaceuticals on heart rate are not accounted for in our estimates of circadian phase, but could be included in future work. Certain types of cardiovascular activity, such as weight lifting and bicycle riding, may also affect heart rate in ways which are not fully accounted for by the way some wearables report activity; use of raw motion data should be investigated.

The relationship between CRHR and other circadian markers raises interesting questions for circadian science. While they generally agree, the data of Timmerman et al. suggests that the CRHR and core-body-temperature (CBT) may in certain cases diverge, although it is unclear how much of this effect was due to masking in their data (Timmerman et al., 1959). Another interesting study shows how high carbohydrate meals may shift CRHR and CBT but not DLMO (Krauchi et al., 2002). Even melatonin onset and offset show different dynamics (Liu and Borjigin, 2005). In animal models simulating jet lag, autonomous circadian clocks in the heart and other peripheral organs shift at different rates than the central clock located in the suprachiasmatic nucleus (SCN) (Damiola et al., 2000; Stokkan et al., 2001). Going forward, to fully understand the human circadian timekeeping system, and its desynchrony when challenged, may require multiple markers.

Easy to use, passive measurements of circadian phase are crucial for the circadian field; our results suggest that these can easily be found in the real-world population via wearable devices. Future work could focus on demographics, as our methods allow for the detailed study how circadian rhythms change with age, location, work schedule and other real-world factors. Together, these results establish CRHR as a practical tool for studying circadian rhythms and their effects on human health.

## Materials and Methods

We estimate CRHR phase using Bayesian uncertainty quantification. To ensure that periods of more frequent measurement do not disproportionately affect results, heart rate and activity data are averaged into five-minute bins. The Fitbit and Mi Band data sets also include sleep timing, which we use to discard all data obtained during sleep (the Apple Watch data set has sleep removed, as the watches were charged overnight.) Data from pairs of consecutive days (centered by a period of sleep) are fit to a 24-hour sinusoidal model with three parameters (phase, mean, amplitude) plus a linear effect from activity using a scaling parameter. Together with two parameters (correlation, noise) describing the autoregressive AR(1) error model, six parameters are sampled from the likelihood using Markov chain Monte Carlo (MCMC). When predicting phase on successive days, the previous day’s fit (plus Gaussian noise with s.d. 1 hour) is used as a prior distribution, and the posterior is sampled instead of the likelihood. Phase estimates (means) and uncertainties are calculated directly from this weighted cloud of fits.

Phase response curves are calculated in 24 one-hour bins. For each day of data, activity is shifted by the CRHR phase estimate from the previous night; total activity within each bin (x) is then compared to the phase difference (y) between the previous night and the following night. The slope within each bin, estimated by simple linear regression, is the average phase shift due to steps during that hour relative to current phase. To avoid overlapping intervals of data, we also tried using phase changes separated by two or more days; as results were similar, we chose to keep the simplest comparison (phase change over 24 hours). Phase response curves are parametrized using a nonlinear least squares fitting of a sinusoidal model with three parameters (phase, mean, amplitude) plus a fourth parameter (trough-to-peak time) allowing the first and second half-periods to have different lengths.

Data sets with a small number of days per subject (e.g., the Apple Watch data set studied in Fig. 2H) do not benefit from the uncertainty propagation of the full Bayesian approach. To reduce bias, we instead use a simple approximation: orthogonalized least squares (Huang et al., 2019a). Heart rates are averaged into one-hour bins; the sinusoidal model and parameters are otherwise identical to the full Bayesian approach, including removing an additive linear effect due to activity. Orthogonalization is necessary to avoid bias due to gaps in data (e.g., during sleep or when watches are charged). Point estimates of CRHR phase are extracted from the least squares fits.

To collect DLMO data for comparison with CRHR, salivary collection began 6 hours prior to each participants’ habitual bedtime; samples were collected via the salivette system every 30 minutes until their bedtime (13 samples total). Participants were in in dim light (<< 50 lux) during the procedure and remained seated at least 10 minutes before each sample. In the event that they had to leave the room, participants wore Uvex glasses to block blue spectrum light. Immediately prior to each scheduled sample collection, participants brushed their teeth with water if they ate or drank anything other than water between samples. Participants provided saliva by chewing on the swab until it became impregnated with saliva and deposited the swab in the appropriate salivette. The samples were centrifuged for two minutes, frozen (−20°C), and assayed using radioimmunoassay. Other aspects of this protocol are as described in (Walch et al., 2019).

